# Bacterial degradation of ctenophore *Mnemiopsis leidyi* organic matter

**DOI:** 10.1101/2023.08.14.553244

**Authors:** Eduard Fadeev, Jennifer H Hennenfeind, Chie Amano, Zihao Zhao, Katja Klun, Gerhard J Herndl, Tinkara Tinta

## Abstract

Blooms of gelatinous zooplankton, an important source of protein-rich biomass in coastal waters, often collapse rapidly, releasing large amounts of labile detrital organic matter (OM) into the surrounding water. Although these blooms have the potential to cause major perturbations in the marine ecosystem, their effects on the microbial community and hence on the biogeochemical cycles have yet to be elucidated. We conducted microcosm experiments simulating the scenario experienced by coastal bacterial communities after the decay of a ctenophore (*Mnemiopsis leidyi*) bloom in the northern Adriatic Sea. Within 24 h, a rapid response of bacterial communities to the *M. leidyi* OM was observed, characterized by elevated bacterial biomass production and respiration rates. Compared to our previous microcosm study of jellyfish (*Aurelia aurita s*.*l*.), *M. leidyi* OM degradation revealed that, despite the fundamental genetic and biochemical differences between the two species, a similar pattern in the bacterial community response was observed. Combined metagenomic and metaproteomic analysis indicated that the degradation activity was mainly performed by *Pseudoalteromonas*, producing a large amount of proteolytic exoenzymes and exhibiting high metabolic activity. Interestingly, the reconstructed metagenome-assembled genome (MAG) of *Pseudoalteromonas phenolica* was almost identical (average nucleotide identity >99%) to the MAG previously reconstructed in our *A. aurita* microcosm study. Taken together our data suggest that blooms of different gelatinous zooplankton are likely triggering a consistent response from natural bacterial communities, with specific bacterial lineages driving the remineralization of the gelatinous OM.

**Importance:** Jellyfish blooms are increasingly becoming a recurring seasonal event in marine ecosystems, characterised by a rapid build-up of gelatinous biomass that collapses rapidly. Although these blooms have the potential to cause major perturbations, their impact on marine microbial communities is largely unknown. We conducted an incubation experiment simulating a bloom of the ctenophore *Mnemiopsis leidyi* in the Northern Adriatic, where we investigated the bacterial response to the gelatinous biomass. We found that the bacterial communities actively degraded the gelatinous organic matter, and overall showed a striking similarity to the dynamics previously observed after a simulated bloom of the jellyfish *Aurelia aurita s*.*l*. In both cases we found that a single bacterial species, *Pseudoalteromonas phenolica*, was responsible for most of the degradation activity. This suggests that blooms of different jellyfish are likely to trigger a consistent response from natural bacterial communities, with specific bacterial species driving the remineralisation of gelatinous biomass.

## Introduction

Blooms of gelatinous zooplankton are increasingly recognized as an important source of organic matter for marine ecosystems worldwide, with possible links to anthropogenic pressures and climate change (Purcell, 2012). Gelatinous organisms have existed for > 600 million years and some even speculate that we may be facing a gelatinous future due to the adaptability of these ancient organisms (Richardson et al., 2009). Particularly coastal environments are more frequently experiencing recurring seasonal bloom events of native or invasive gelatinous zooplankton species (i.e., the cnidarian subphylum Medusozoans and the phylum Ctenophora, hereinafter collectively coined jellyfish). Jellyfish blooms are characterized by a rapid buildup of biomass up to ∼ 6 mg C m^−3^, and sometimes, particularly in coastal areas, even exceeding ∼ 10 g C m^−3^ (Lucas et al., 2014; Luo et al., 2020). These blooms are often short-lived (weeks to month) followed by a sudden collapse of the population, leading to a massive release of detrital organic material (OM) into the ecosystem (Pitt et al., 2014). The released detrital OM has a high protein content and is typically characterized by low carbon (C) to nitrogen (N) molar ratios (C:N=∼4.5:1), in contrast to the carbon-rich OM released by phytoplankton (C:N=6.6:1) (Pitt et al., 2009). Jellyfish-derived OM (jelly-OM) is more accessible to heterotrophic microbes than crustacean zooplankton due to the lack of a chitinous exoskeleton. About half of the released jelly-OM is in the dissolved phase (< 0.8 µm) and thus, is exclusively accessible to marine pelagic bacteria that respond to it within hours (Tinta et al., 2020). The fraction of the jelly-OM not re-mineralized by bacteria in the water column sinks to the bottom, where it is degraded by benthic bacterial communities (Tinta et al., 2021 and the reference therein). As marine bacteria are the key drivers of oceanic biogeochemistry (Azam, 1998), unraveling their role in the re-mineralization of gelatinous OM is important for further understanding the impact of jellyfish blooms on coastal marine environments and to properly incorporate bacteria-jellyfish interactions into oceanic carbon budgets and biogeochemical cycles (Tinta et al., 2021).

A rapid increase in OM availability during phytoplankton blooms has been shown to trigger a network of metabolic processes within marine bacterial communities (Teeling et al., 2016, 2012). However, due to the temporal and spatial patchiness of jellyfish blooms, *in situ* observations of bacteria-jelly OM interactions are scarce and most of the existing knowledge is derived from incubation experiments. Previous studies have shown that bacterial communities thrive on jelly-OM, reaching considerably higher growth rates than otherwise reported for pelagic marine bacteria (Blanchet et al., 2015; Tinta et al., 2021; Titelman et al., 2006). Our previous microcosm experiments focusing on microbial degradation of jelly-OM from the cosmopolitan bloom-forming scyphozoan jellyfish *Aurelia aurita s*.*l*. revealed that natural marine bacterial communities are capable of consuming most of its detrital OM within 1.5 days (Tinta et al., 2020). We found that during this short time period the bacterial community incorporated a large fraction of the introduced organic carbon into its biomass (indicated by a bacterial growth efficiency of 65 ± 27%). Furthermore, the bacterial communities consumed more than 97% of the dissolved proteins and 70% of the dissolved amino acids of the initial jelly-OM pool (Tinta et al., 2020).

Bioavailability of gelatinous detrital OM favors specific bacterial lineages and results in compositional changes of the communities (Blanchet et al., 2015; Dinasquet et al., 2013; Tinta et al., 2012). We identified that during *A. aurita* OM degradation, the taxonomic lineages *Pseudoalteromonas, Alteromonas*, and *Vibrio* (all within Gammaproteobacteria) were the most metabolically active members of the bacterial community (Tinta et al., 2022). Using a proteomics approach, we have found that *Pseudoalteromonas* played an important role in the initial extracellular degradation of the protein-containing OM, *Alteromonas* processed carbohydrates and organophosphorus compounds, and *Vibrio* exhibited a cheater lifestyle exploiting the by-/end-products of the metabolic processes carried out by the other two lineages (Tinta et al., 2022). Collectively, these observations suggest that the introduction of gelatinous OM can affect natural marine bacterial communities by stimulating the growth of specific bacterial taxa, as well as promoting specific degradation pathways and complex metabolic interactions within the communities.

The term ‘jellyfish’ is not restricted to “true jellyfish” (i.e., class Scyphozoa) and often refers to other gelatinous zooplankton, such as comb jellies (phylum Ctenophora), which also form massive blooms. Some jellyfish and ctenophore species are also invasive and ever since the early 1980s, when the lobed ctenophore *Mnemiopsis leidyi*, native to the western Atlantic Ocean, invaded the Black Sea, its blooms have become an increasingly common phenomenon in Eurasian seas (Ghabooli et al., 2013; Shiganova et al., 2019). Since 2016, large-scale blooms of *M. leidyi* are recurring annually from mid-summer to late autumn in the northern Adriatic, and specifically in the Gulf of Trieste (Malej et al., 2017). Few previous studies on the interaction between pelagic bacterial communities and *M. leidyi* blooms have shown that, similar to other jellyfish, ctenophore detrital OM stimulates bacterial growth and causes a shift in the bacterial communities towards the dominance of Gammaproteobacteria (Dinasquet et al., 2012). Despite these similarities, differences in life history traits, genetic background and hence distinct physiological and biochemical characteristics (e.g., lack of toxins in ctenophores) suggest that the composition of jellyfish and comb jellyfish detrital OM is not the same. We hypothesized that the differences in the OM might lead to different responses in natural bacterial communities. To test that, we replicated previously conducted microcosm experiments on the degradation of *A. aurita* using *M. leidyi* instead. We then characterized the bacterial metabolic response to ctenophore detrital OM and compared the results with those obtained for *A. aurita*.

## Results

The design of the microcosm experiment aimed at mimicking the ecological scenario experienced by pelagic bacterial communities in the Adriatic Sea during a decay of *M. leidyi* bloom and the subsequent introduction of ctenophore detrital OM. The experiment consisted of six microcosms, inoculated with the same ambient seawater and hence the same microbial community. Three of the microcosms were treated with ctenophore detrital OM (further termed “Cteno-OM”, see Materials and Methods section for details on pre-processing of ctenophores) and the other three without amendment served as a control.

Bacterial communities in the Cteno-OM microcosms reached up to one order of magnitude higher cell abundances than the control treatments (Figure 1A). Bacterial cells abundance peaked after about 21 h in all microcosms, except in one of the Cteno-OM replicates where bacterial abundances continued to increase for a longer time (58 h). During the exponential phase (9-21 h), growth rates differed between the treatments, reaching 0.16-0.21 cells h^-1^ in the Cteno-OM microcosms compared to 0.02-0.10 cells h^-1^ in the control. Based on the changes in dissolved organic carbon (DOC) concentrations and the increase in bacterial biomass (Figure 2), the estimated total bacterial production during the exponential phase was 7-10 µg C L^-1^ h^-1^ and 0.2-0.8 µg C L^-1^ h^-1^ in the Cteno-OM and control microcosms, respectively. Furthermore, bacterial growth efficiency (BGE), calculated from the increase in the bacterial abundance converted into biomass production and the decrease in the concentration of DOC, reached 18-27% in the Cteno-OM (n=2) compared to 0-5% in the control microcosms (n=2). Specific gammaproteobacterial taxa (*Alteromonadales, Pseudoalteromonadales*, and *Vibrionales*) reached two to three orders of magnitude higher cell specific productivity in the Cteno-OM than in the control treatments (see Materials and Methods section for details) (Figure 1B). Interestingly, at the peak of the bacterial abundance, *Pseudoalteromonadales* comprised 20-75% of the total cell-specific bacterial productivity. Cell-specific bacterial respiration showed overall similar trends as production (Figure 1C).

**Figure 1.**
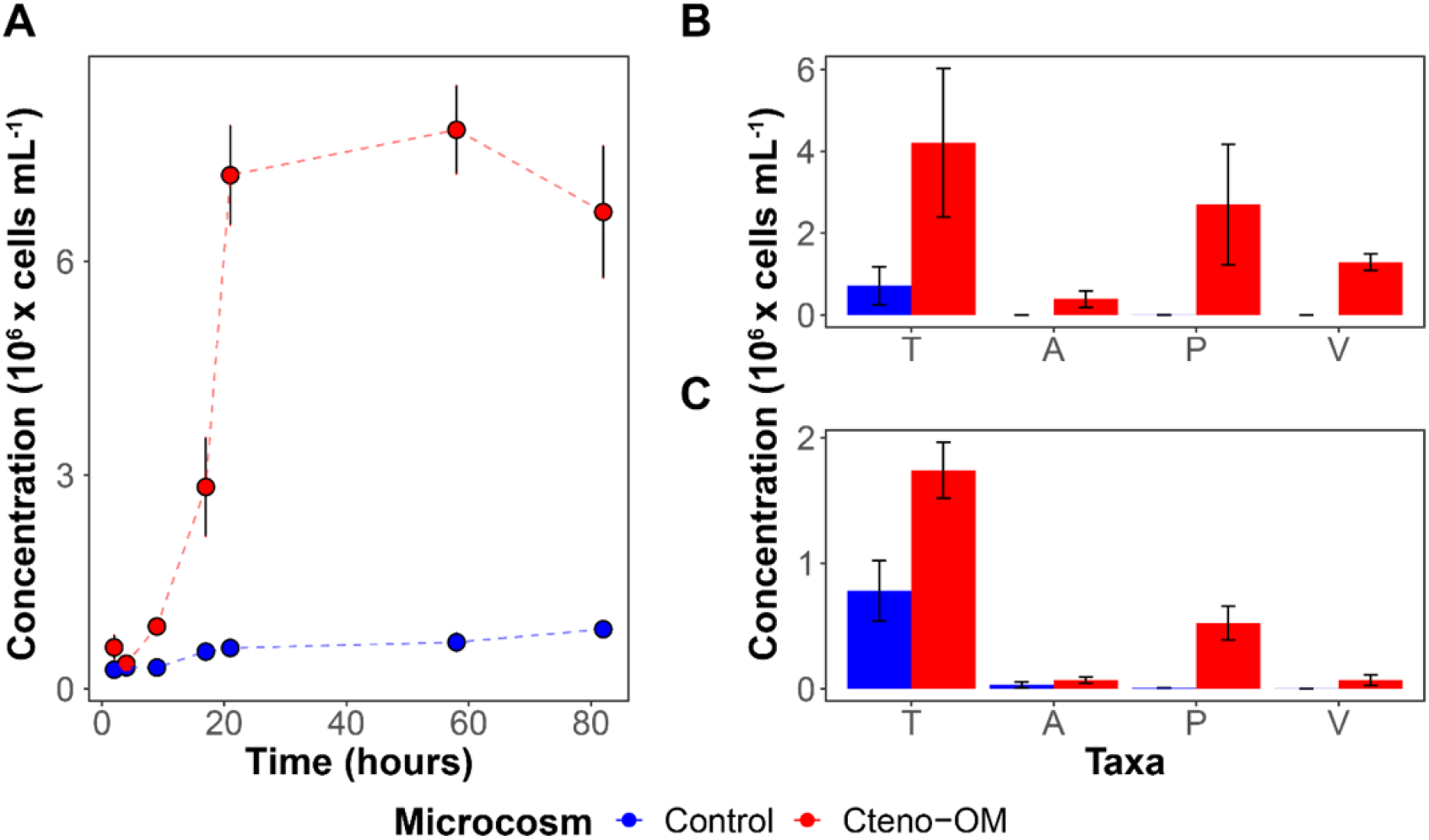
Abundance and activity of bacterial communities in the microcosms. **(A)** Total bacterial abundances in Cteno-OM (red) and control (blue) microcosms estimated by DAPI counts. Mean abundance of HPG incorporating **(B)** and respiring **(C)** bacterial cells at the peak of the abundance estimated using targeted fluorescence in-situ hybridization (FISH) probes. In panels B and C: T-all bacterial cells, A - *Alteromonas*, P - *Pseudoalteromonas*, V - *Vibrio*. Error bars represent standard errors between biological replicates.

**Figure 2.**
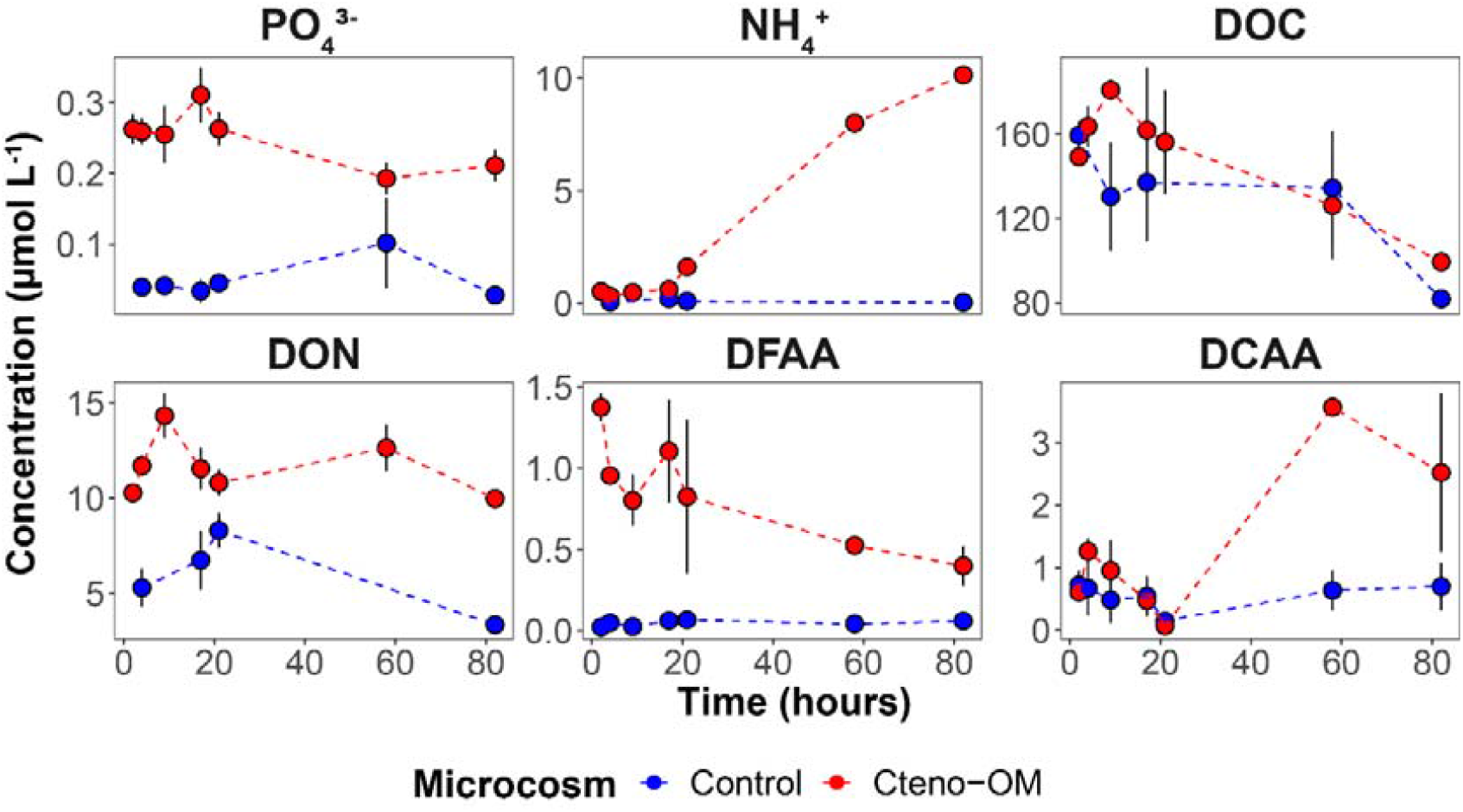
Nutrient concentrations in the microcosms over time. PO_4_^3-^ - phosphate, NH_4_^+^ - ammonium, DOC - dissolved organic carbon, DON - dissolved organic nitrogen, DFAA - dissolved free amino acids, DCAA - dissolved combined amino acids. Error bars represent standard error between biological replicates.

The exponential growth of the bacterial communities in the Cteno-OM microcosms was associated with decreasing orthophosphate (PO_4_ ^3-^) concentrations (Figure 2). The increase in total dissolved nitrogen (TDN) concentration in the Cteno-OM microcosms over time was mainly due to a 20-fold increase in ammonium (NH_4_^+^) from 0.54±0.15 to 10.15±0.15 µmol L^-1^, while concentrations of NO_3_^-^ and NO_2_ ^-^ remained constant (0.06±0.01 and 7.84±0.01 µmol L^-1^, respectively). The initial concentration of dissolved free amino acids (DFAA) was 100 times higher in the Cteno-OM microcosms than in the control and decreased during the entire incubation period from 1.37±0.08 to 0.40±0.12 µmol L^-1^ (Figure 2). Concentrations of the dissolved combined amino acids (DCAA) in Cteno-OM microcosms increased during the first 58 h from 0.61±0.06 to 3.57±0.14 µmol L^-1^. Both, DFAA and DCAA concentrations remained essentially constant in the control microcosms throughout the experiment (DFAA: from 0.02±0.01 to 0.06±0.01 µmol L^-1^, DCAA: from 0.73±0.22 to 0.70±0.37 µmol L^-1^).

Using mass spectrometry, we compared the number of identified *M. leidyi* proteins in the starting Cteno-OM dry material and the Cteno-OM amended microcosms at the peak of bacterial abundance. We found that of the 6128 *M. leidyi* proteins detected in the Cteno-OM dry material, only half (2768-4543) were still detected in the Cteno-OM microcosms at the peak of bacterial abundance (after 21 h; Figure S1). We then analyzed the proteins associated with bacteria in both, the cellular (> 0.22 µm) and extracellular (< 0.22 µm) fraction of each microcosm, as well as in the unamended controls. The cellular fraction of the inoculum contained approximately one third more bacterial proteins (total of 6256) than the Cteno-OM (2170-6047) and the control (3088-4869) microcosms (Figure S2; Table S1). In contrast, in the extracellular fraction, no major differences were observed in the total number of identified proteins (inoculum: 3823, Cteno-OM: 3310-4975, control: 3735-4026). A total of 461 proteins were observed across the entire dataset (i.e., in both fractions of the Cteno-OM and control mesocosms, as well as the unamended seawater), 867 proteins were shared between all cellular fractions, and 1170 proteins were shared between all the extracellular fractions. In each microcosm, half of the proteins were observed in both size fractions. However, the composition of proteins was significantly different between the fractions (PERMANOVA test; F_1,13_=1.73, R^2^=0.11, p=0.02; Figure 3), and in both fractions, the composition of proteins was significantly different between the treatments (PERMANOVA test; F_2,13_=1.49, R^2^=0.19, p=0.02).

**Figure 3.**
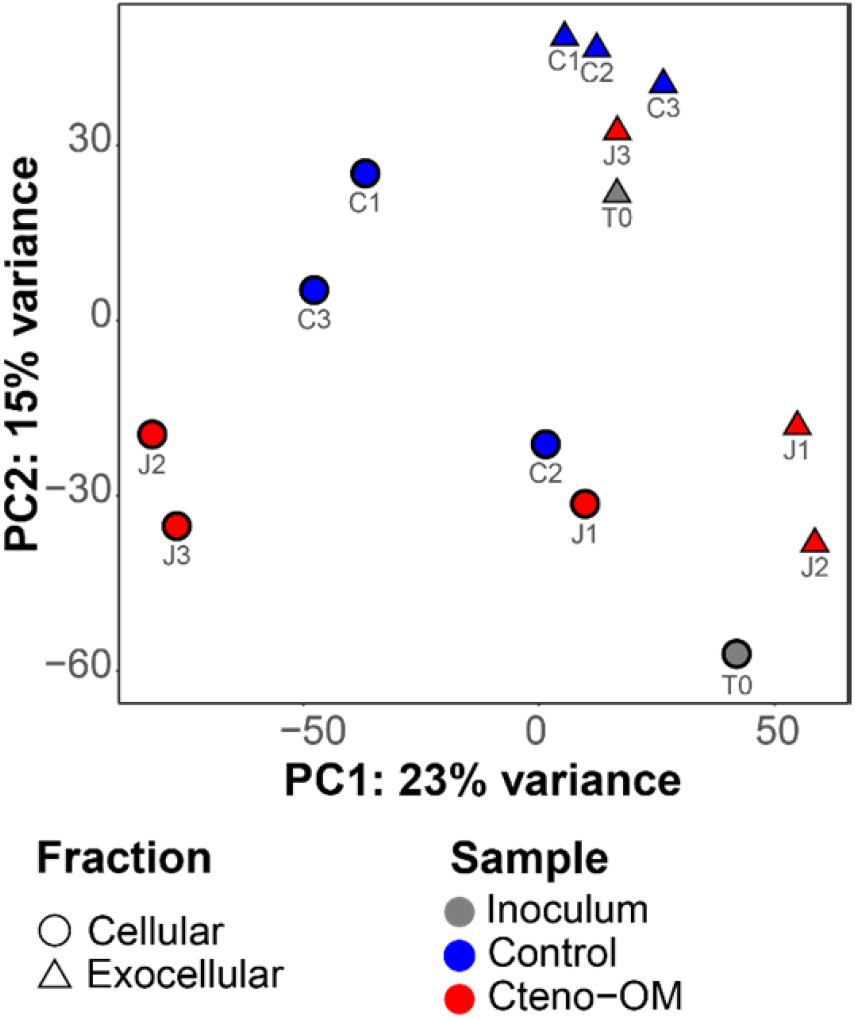
Dissimilarity of the protein composition in the microcosms and the inoculum bacterial communities. Principal component analysis (PCA) was performed on variance stabilized protein composition in both, the cellular (circle) and extracellular (triangle) fraction.

The taxonomic origin of the identified proteins strongly differed between the bacterial inoculum and the community in the microcosms (Figure 4). The inoculum community contained mostly proteins from the class Alphaproteobacteria (57% of cellular and 34% of extracellular proteins), followed by Gammaproteobacteria (21% of cellular and 53% of extracellular proteins). In both Cteno-OM and the control microcosms, however, most of the proteins originated from lineages in the Gammaproteobacteria (71-80% of cellular and 63-71% of extracellular proteins), while Alphaproteobacteria comprised only 16-21% of cellular and 23-26% of extracellular proteins at the peak of bacterial abundance. In both fractions, the order *Alteromonadales* (class Gammaproteobacteria) contributed to the cellular fraction 52-63% of the total proteins in the Cteno-OM microcosms and 28-43% of the total proteins in the control, while no significant difference was observed in the extracellular fraction between the Cteno-OM and the control treatment (28-37% and 22-34% of the proteins, respectively). In the control microcosms, the order *Oceanospirillales* (class Gammaproteobacteria) exhibited a higher relative abundance of proteins than in the Cteno-OM treatment (16-23% vs. 9-12% of the proteins in the cellular fraction, 28-37% and 22-34% of the proteins in the extracellular fraction). The order *Pelagibacterales*, comprising the largest portion of proteins among Alphaproteobacteria did not show differences in the relative abundance of proteins between the two treatments, comprising 4-26% of the proteins in the cellular fraction and 8-25% of the proteins in the extracellular fraction.

**Figure 4.**
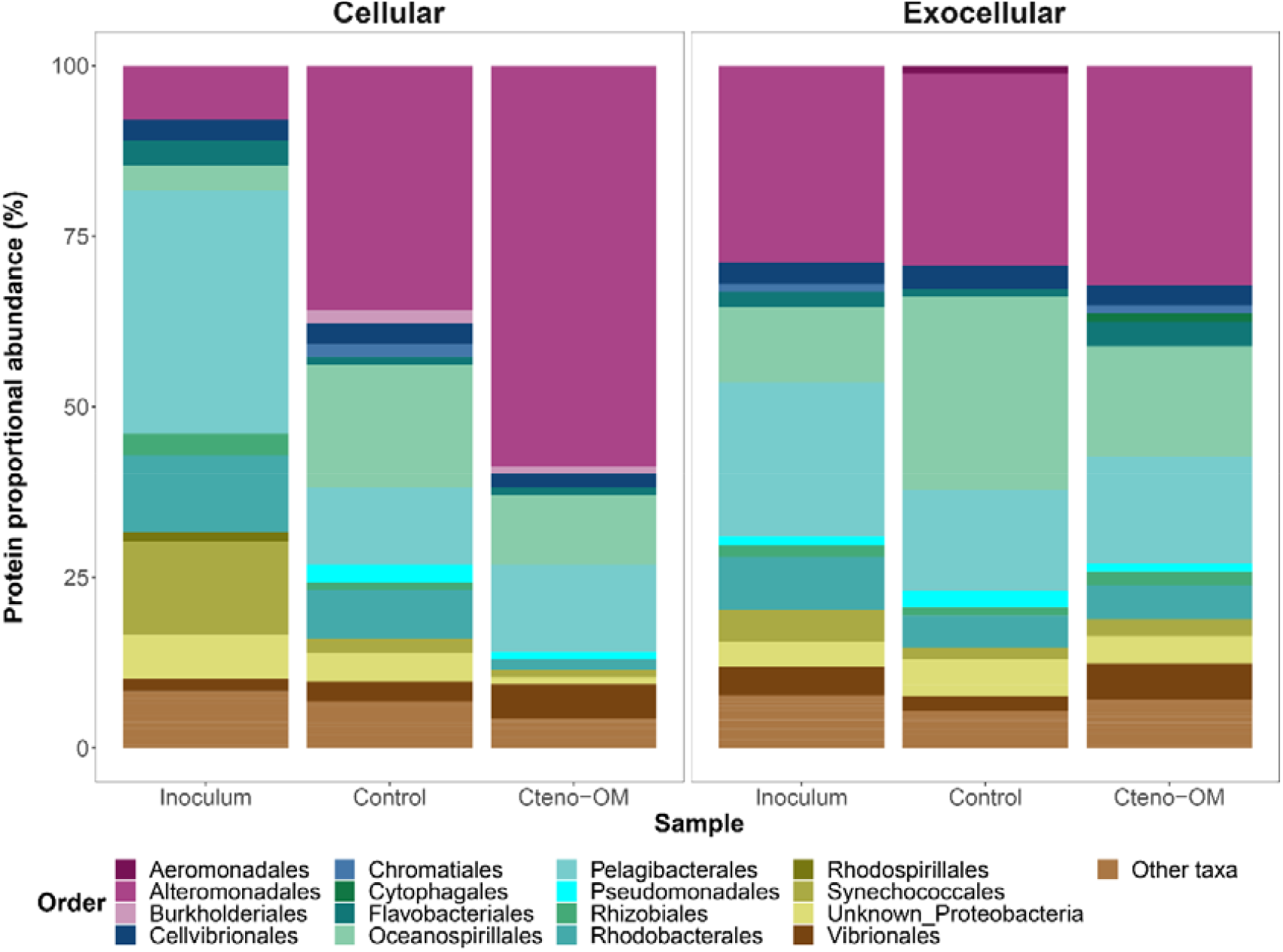
Taxonomic breakdown of bacterial proteins detected in the cellular and exocellular fractions. The stack bar for the two treatments (Cteno-OM and unamended control) represents mean value of three replicates.

To identify significantly enriched proteins between the Cteno-OM and the control microcosms, we implemented the DESeq2 enrichment algorithm on both the cellular and the extracellular protein datasets (Figure 5). We identified 490 and 603 proteins in the cellular and extracellular fraction, respectively, which were significantly enriched in the Cteno-OM microcosms (adjusted *p* value < 0.1; Table S2). In both fractions combined, the bacterial genus with the largest number of enriched proteins was *Pseudoalteromonas* (365 proteins; order *Alteromonadales*). The enriched proteins of this genus were mostly associated with the Clusters of Orthologous Genes (COG) categories J - Translation, ribosomal structure and biogenesis (60 proteins, mostly various tRNA synthetases and other ribosomal proteins), E - Amino acid transport and metabolism (40 proteins, among them Leucyl-, Xaa-Pro-, and N-aminopeptidases), O - Posttranslational modification and protein turnover (32 proteins, among them Lon and Serine proteases, as well as various peptidases). The genus *Alteromonas* also contained a large number of significantly enriched proteins (48 proteins). Enriched proteins of *Alteromonas* were associated to COG categories O - Posttranslational modification and protein turnover (7 out of the total 8 proteins were ATP-dependent Lon protease) and J - Translation, ribosomal structure and biogenesis (7 proteins, containing various tRNA synthetases and other ribosomal proteins). The second largest group of significantly enriched proteins in the Cteno-OM microcosms was affiliated to the genus *Vibrio* (61 proteins; order *Vibrionales*) with the enriched proteins associated to COG categories G - Carbohydrate transport and metabolism (12 proteins of various metabolic functions), J - Translation, ribosomal structure and biogenesis (11 mostly ribosomal proteins), and E - Amino acid transport and metabolism (8 proteins, among them various proteins associated with ABC-type transport systems). In the control microcosms, we found 302 and 339 significantly enriched proteins in the cellular and extracellular fractions, respectively (Table S2). In both fractions, the enriched proteins were mostly associated with various genera within the orders *Alteromonadales* and *Pelagibacterales*, with no clear trends in their COG affiliations.

**Figure 5.**
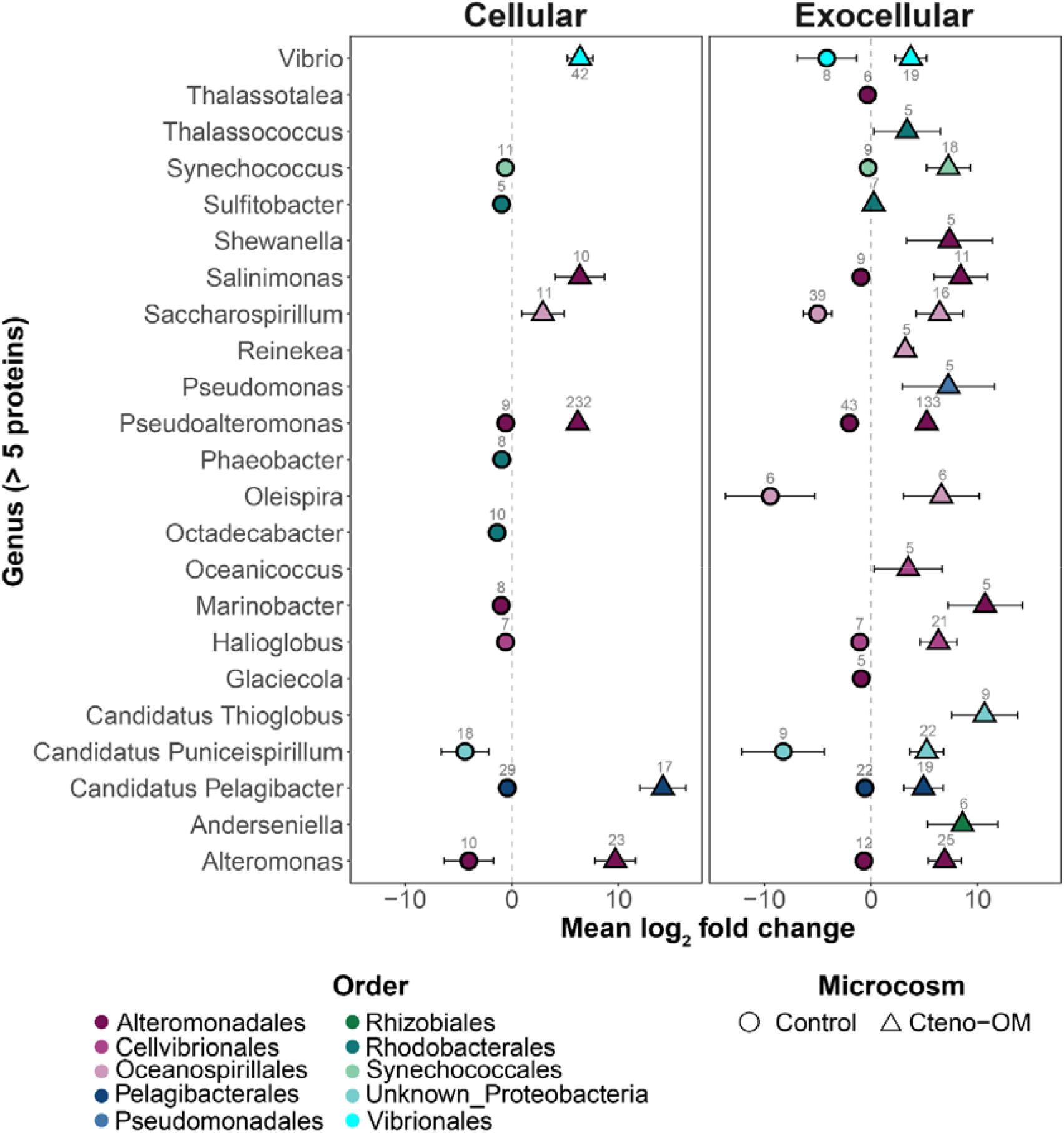
Differential abundance of proteins according to taxonomic origins between Cteno-OM and control microcosms. The mean log_2_ fold-change of significantly enriched proteins was calculated according to grouping of distinct genera and fraction (cellular and extracellular). Positive log_2_ fold-change values represent enrichment in Cteno-OM microcosms, and negative values represent enrichment in control microcosms.

Using a co-assembled metagenome of DNA libraries from both the Cteno-OM and the control microcosms as well as the inoculum seawater sample (described in Materials and Methods), we reconstructed 33 bacterial metagenome-assembled genomes (MAGs), all larger than 1.2 Mbp, with estimated completeness of >75%, and redundancy <8% (Figure 6). The coverage of the MAGs by reads from each library (i.e., different samples) revealed that the DNA of two MAGs was notably more abundant in the Cteno-OM than in the control treatment. These two MAGs were taxonomically affiliated to the bacterial species *Marinobacterium jannaschii* and *Pseudoalteromonas phenolica* (Figure S3). The reconstructed *P. phenolica* MAG (Bin_84) had the highest read coverage among all MAGs and comprised 506 contigs with a total of 4145 coding genes (overall length 4.2 Mbp) and was highly similar to the previously reconstructed MAG in the *A. aurita s*.*l*. degradation experiment (average nucleotide identity - 99%)(Tinta et al., 2022). The reconstructed *P. phenolica* MAG shared 1143 genes with all other *P. phenolica* genomes (i.e., core genes) and contained 122 genes not present in any other genome (Figure S4 and Table S3).

**Figure 6.**
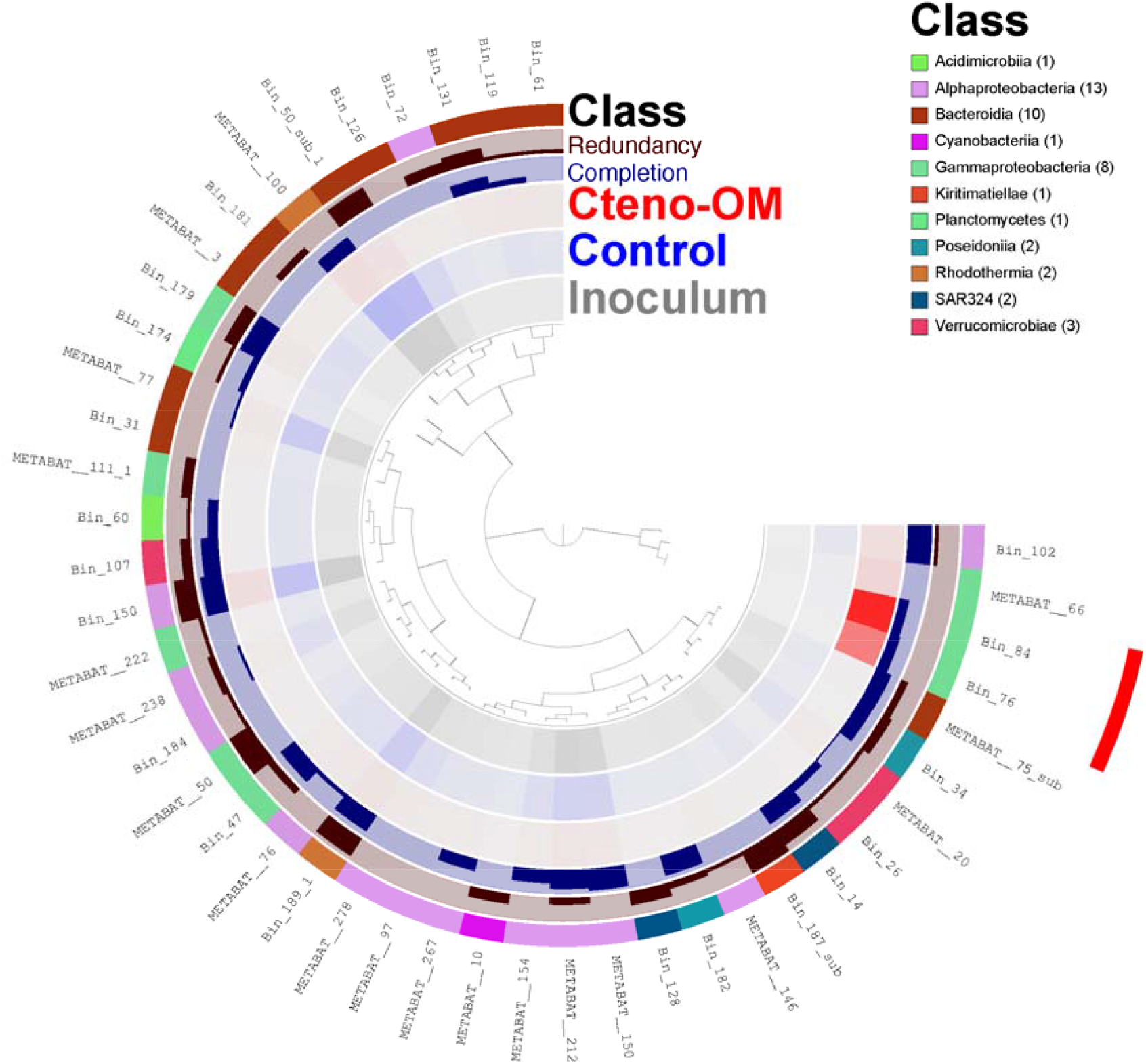
Reconstructed MAGs from the co-assembled metagenome. The color density in the three inner rings represents the normalized average coverage of each MAG by reads from each individual metagenome (Cteno-OM, control, and inoculum). The MAGs were clustered using Euclidian distance between their read coverages. The MAGs marked in red had higher read coverage in the Cteno-OM metagenome. Completion represents the estimated completeness of the MAG ranging from 75-100%. The redundancy represents the estimated sequence contamination in the MAG ranging from 0-8%.

Of all the MAGs reconstructed, the *P. phenolica* MAG was associated with the largest number of proteins in the microcosms (194-505 in the Cteno-OM and 209-287 in the control treatments). In the Cteno-OM treatments, these proteins accounted for 21-27% in the cellular and 7-8% in the extracellular fraction of all proteins in the dataset (Figure 7). To characterize the potentially enriched metabolism of *P. phenolica* in the Cteno-OM treatments, we reconstructed its metabolic pathways using the genomic information in the MAG and combined it with the identified significantly enriched proteins. Among those *P. phenolica* MAG-associated proteins, 149 proteins were significantly enriched in the Cteno-OM treatments (in contrast to only 16 significantly enriched proteins in the control treatments). The genomic information of the *P. phenolica* MAG contained numerous complete metabolic pathways (completeness defined by the presence of >70% of the required genes; Table S3). We found significantly enriched proteins associated with glycolysis (K01803 - EC:5.3.1.1) and gluconeogenesis (K01610 – EC:4.1.1.49) pathways, as well as with pyruvate oxidation (K00382 – EC:1.8.1.4). Among the enriched proteins we also found two subunits (alpha - K02111 and beta - K02112) of F-type ATPase (EC:7.1.2.2) and one subunit (iron-sulfur subunit - K00411) of the Cytochrome c reductase (EC:7.1.1.8), both associated with oxidative phosphorylation. These were complemented by enrichment of two different proteins (K03183 - EC:2.1.1.201 and K00568 - EC:2.1.1.222) in the ubiquinone biosynthesis pathway, a key redox cofactor of the electron transfer chain. In addition, we found enrichment of the acyl-CoA dehydrogenase (K00249 - EC:1.3.8.7) playing a key role in beta-oxidation of fatty acids. However, this enzyme is also involved in the leucine degradation pathway, in which we also observed an enrichment of leucine dehydrogenase (K00263 - EC:1.4.1.9) and dihydrolipoamide dehydrogenase (K00382 - EC:1.8.1.4). Also, enriched proteins such as glutamate dehydrogenase (K00260 - EC:1.4.1.2) are potentially linked to the degradation of other amino acids, such as alanine, arginine and of taurine. We further observed enrichment of various proteins associated with protein metabolism, such as the protein export system SecY/Sec61 (K03070 - EC:7.4.2.8) and signal peptidases (K03100 – EC:3.4.21.89).

**Figure 7.**
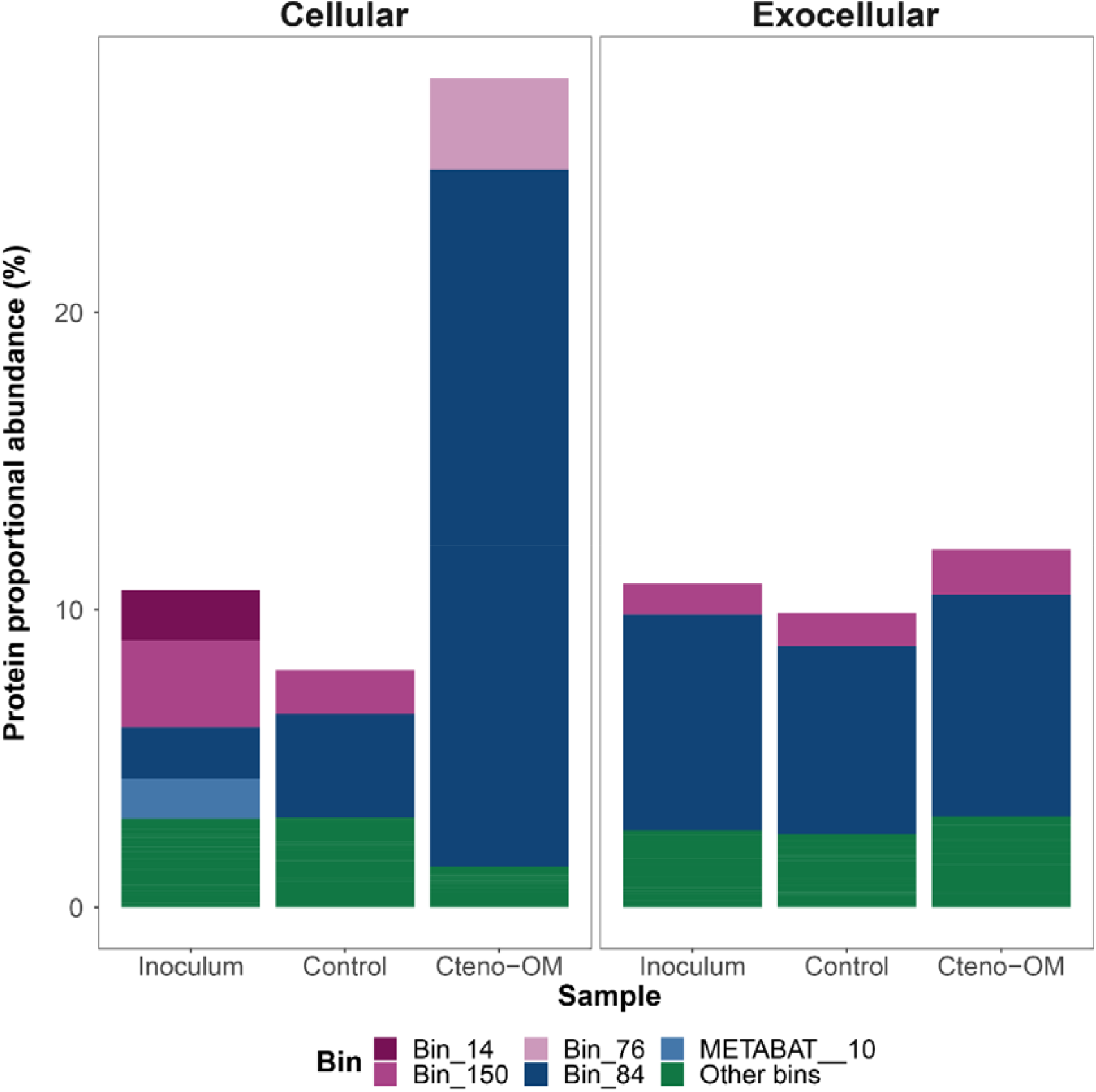
Proportional abundance of proteins associated with MAGs. The MAGs with a total protein proportion below 1% were grouped under ‘other bins’. Bin_14 – uncl. SAR324 clade; Bin_150 - uncl. *Rhodobacteraceae*; Bin_76 – *M. jannaschii*; Bin_84 - *P. phenolica*; METABAT__10 uncl. *Cyanobiaceae*.

## Discussion

Gelatinous zooplankton includes a group of diverse organisms that share several traits such as planktonic lifestyle and trophic position. These organisms are commonly referred to as ‘jellyfish’ and often lumped together in biogeochemical oceanic models and budgets. However, despite their similarities, these organisms are characterized by different life history traits and are genetically, and therefore physiologically and biochemically distinct. Comparing the biochemical composition of the ctenophore *M. leidyi* OM with that of the scyphozoan jellyfish *A. aurita s*.*l*., we found that the *A. aurita s*.*l*. contains more than twice the amount of dissolved organic carbon, total dissolved nitrogen, and phosphate (Table 1). This raised the question of whether the biochemical differences affect the ‘fate’ of the gelatinous detrital OM. Do marine ambient bacterial communities exhibit the same response to gelatinous OM regardless of its source and distinct biochemical features? Understanding this is crucial for correctly incorporating bacteria-jellyfish interactions into oceanic biogeochemical models.

**Table 1:**
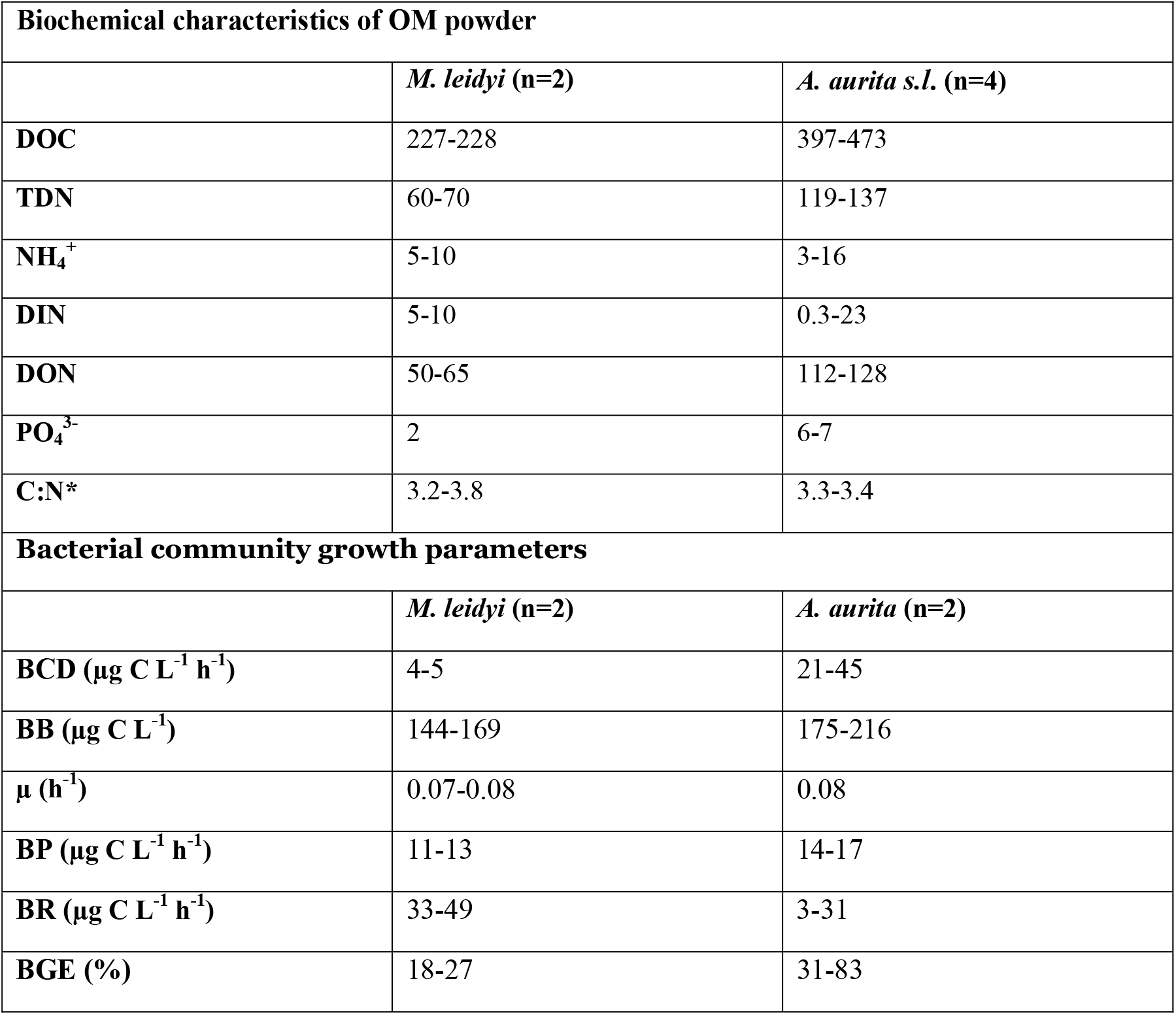
Comparison between *M. leidyi* and *A. aurita s*.*l*. leaching and microcosm experiments. All biochemical values represent concentrations of dissolved analyte (< 0.8μm) in µmol g^-1^ powder (i.e., per dry weight of a subsample of *M. leidyi* or *A. aurita s*.*l*. representative populations). Due to a substantially different community dynamic, one replicate in each experiment was excluded from growth estimates. DOC - dissolved organic carbon; TDN - total dissolved nitrogen; DIN - dissolved inorganic nitrogen = NH_4_^+^+NO_2_^-^+NO_3_^-^; DON - dissolved organic nitrogen = TDN-DIN; BCD - bacterial carbon demand; BB - bacterial biomass; µ- growth rate; BP - bacterial production; BR - bacterial respiration; BGE - bacterial growth efficiency; * note that C:N ratio is calculated for dissolved fraction, i.e., DOC:DON.

| Biochemical characteristics of OM powder |  |  |
| --- | --- | --- |
|  | *M. leidy* (n=2) | *A. aurita s.l.* (n=4) |
| DOC | 227-228 | 397-473 |
| TDN | 60-70 | 119-137 |
| $\text{NH}_4^+$ | 5-10 | 3-16 |
| DIN | 5-10 | 0.3-23 |
| DON | 50-65 | 112-128 |
| $\text{PO}_4^{3-}$ | 2 | 6-7 |
| C:N* | 3.2-3.8 | 3.3-3.4 |
| Bacterial community growth parameters |  |  |
|  | *M. leidy* (n=2) | *A. aurita* (n=2) |
| BCD ($\mu\text{g C L}^{-1} \text{ h}^{-1}$) | 4-5 | 21-45 |
| BB ($\mu\text{g C L}^{-1}$) | 144-169 | 175-216 |
| $\mu$ ($\text{h}^{-1}$) | 0.07-0.08 | 0.08 |
| BP ($\mu\text{g C L}^{-1} \text{ h}^{-1}$) | 11-13 | 14-17 |
| BR ($\mu\text{g C L}^{-1} \text{ h}^{-1}$) | 33-49 | 3-31 |
| BGE (%) | 18-27 | 31-83 |

To start answering these questions, we replicated the experimental design of our previously published study on bacterial degradation of *A. aurita s*.*l*. OM (Tinta et al., 2022), to investigate the response of pelagic bacterial communities to OM of *M. leidyi*. The addition of ctenophore OM to the microcosms promoted rapid bacterial growth and activity, similar to the *A. aurita* treatments (Table 1). However, the biomass reached by the bacterial communities in Cteno-OM microcosms was much lower compared to the *A. aurita s*.*l*. experiment, likely due to the higher DOC content of *A. aurita s*.*l*. OM. Bacterial growth in Cteno-OM microcosms led to a depletion of DOC and phosphate and an accumulation of ammonia and dissolved combined amino acids. The DCAA accumulation most likely represented an enhanced extracellular enzymatic cleavage of proteins and peptides, as well as slower bacterial utilization rates, compared to DFAA (Keil and Kirchman, 1993; Lee, 1993). Additional evidence for protein degradation was the much lower number of *M. leidyi* proteins identified in the Cteno-OM microcosms at the peak of bacterial abundance, as compared to the initial *M. leidyi* OM protein content.

We observed that the introduction of *M. leidyi* detrital OM led to an increased activity of *Vibrionales, Pseudoalteromonadales*, and *Alteromonadales*, the latter is often found enriched in natural seawater communities during phytoplankton blooms (Heins et al., 2021; Teeling et al., 2016, 2012). At the peak of bacterial abundance, the order *Alteromonadales* accounted for more than half of the bacterial proteins identified in the Cteno-OM microcosms. Protein enrichment analysis between the Cteno-OM and the control microcosms revealed that most of the enriched proteins originated from the genera *Pseudoalteromonas* and *Alteromonas*, and were linked to protein hydrolysis, as well as translational and ribosomal processes (t-RNAs synthetases). To elucidate the specific metabolic pathways these enriched proteins were involved in, we reconstructed a *P. phenolica* MAG, which dominated the bacterial community at the peak of bacterial abundance. The reconstructed *P. phenolica* MAG was genomically identical to an *Pseudoalteromonas* MAG acquired from the *A. aurita s*.*l*. microcosm experiment (Tinta et al., 2022).This bacterial species has been previously characterized as a major producer of proteases, lipases, and other hydrolytic enzymes (Chen et al., 2020; Johnson et al., 2017). In our study, we found that *P. phenolica* was actively involved in the hydrolysis of the protein-rich ctenophore OM through the production of a wide range of proteases. The hydrolysis products (i.e., amino acids) were then taken up by *P. phenolica* cells and further degraded through various amino acid metabolic pathways (e.g., leucine and taurine). We also found evidence for lipid oxidation associated with the ctenophore OM (e.g., acyl-CoA dehydrogenase). The products of these degradation pathways have fueled the central metabolism of *P. phenolica* and thereby, elevating its activity, which most likely led to the high abundance of metabolically active *Pseudoalteromonas* as observed by our epifluorescence microscopy-based approaches (Figure 1B-C).

The *M. leidyi* and *A. aurita s*.*l*. microcosm experiments used ambient seawater from the same geographical location (same sampling station), but one year apart. The experiments showed that the magnitude of the bacterial response to the addition of gelatinous OM varied according to its biochemical composition and concentration of specific compounds (Table 1). However, the striking similarity between the key bacterial players in both experiments, and the consistency with previous jellyfish incubation experiments (Condon et al., 2011; Dinasquet et al., 2012; Tinta et al., 2020, 2012) suggests that copiotrophic bacterial lineages, such as *P. phenolica*, drive a fairly consistent metabolic response of marine bacterial communities to natural blooms of different gelatinous organisms. Therefore, we postulate that in order to assess the biogeochemical impact of ‘jellyfish’ blooms, special attention must be paid to the identification of bacterial lineages with specific enzymatic and metabolic capabilities for the remineralization of gelatinous OM.

## Materials and methods

### Preparation of *M. leidyi* OM powder and its biochemical characterization

A total of 21 individuals of *M. leidyi* were collected in the Gulf of Trieste, northern Adriatic Sea, during the summer bloom in August 2019. The ctenophores were collected on different days and at different locations to account for potential spatial and temporal heterogeneity in the bloom population. Individuals were sampled from the surface of the water column using a large acid-cleaned plastic bucket and stored in zip-lock bags at -20°C within 1 h. Each ctenophore was then freeze-dried at - 45°C for 7 d and the dry material of all ctenophores was pooled and homogenized with a sterilized pestle and agate mortar. The Cteno-OM powder was then stored in acid- and Milli-Q water-rinsed and combusted glass vials at -20°C. To minimize the risk of contamination and degradation of the Cteno-OM powder, care was taken to work under sterile conditions and on ice at all intermediate steps.

To determine its chemical composition leaching experiments were conducted as follows: 250 mg of the Cteno-OM powder was dissolved in 1 L of artificial seawater (prepared according to (Kester et al., 1967) in an acid- and Milli Q water-rinsed and combusted glass Erlenmeyer flask. To ensure maximized dissolution of the powder, the Erlenmeyer flask was placed on a shaker in the dark at room temperature for 24 h. From each flask a technical duplicate was collected for biochemical characterization as described below. In addition, the absence of possible bacterial contamination was checked by measuring bacterial abundance.

### Experimental design of the microcosms

In August 2019, 60 L of seawater were collected from 5 m depth in the center of the Gulf of Trieste and aged in acid-washed and Milli-Q water-rinsed 20 L Nalgene carboys (ThermoFisher Scientific, Rockford, IL, USA) for about one month at room temperature in the dark, and then filtered through an 0.22 µm polycarbonate filter to remove microbial cells. In September 2019, an additional water sample was collected at the same sampling location and pre-filtered through a 1.2 µm polycarbonate filter to remove most of the non-bacterial particulate matter from the sample. Then, the aged and the ‘fresh’ seawater were distributed into six 10 L borosilicate glass bottles in a 9:1 ratio. Based on an average of ca. 100 *M. leidyi* specimens m^-3^ observed during sampling and their dry weight of ca. 1 g each, a total of 1 g of Cteno-OM powder was added to three microcosms, to reach final concentration of 100 mg L^-1^. Three other microcosms, with no Cteno-OM amendment served as control. All bottles were incubated in the dark at the ambient seawater temperature measured during sampling (∼ 24°C) and mixed gently prior to subsampling.

### Dissolved organic carbon and nitrogen

Samples for dissolved organic carbon (DOC) and total dissolved nitrogen (TDN) were filtered through combusted GF/F filters (Whatman plc, Maidstone, UK) using acid-washed, Milli-Q water-rinsed, and combusted glass filtration system. Approximately 30 mL of the filtrate was collected into acid-, Milli-Q water rinsed and combusted glass vials and acidified with 12 M HCl (∼100 µL per ∼ 20 mL of sample) to reach a final pH < 2 and stored at 4°C until analysis. Both DOC and TDN analyses were performed by a high temperature catalytic method using a Shimadzu TOC-L analyser (Shimadzu, Kyoto, Japan) equipped with a total nitrogen unit (Hansell et al., 1993). The calibration for non-purgeable organic carbon was done with potassium phthalate and for TDN potassium nitrate was used. The results were validated with Deep-Sea Reference (DSR) water for DOC and TDN (CRM Program, Hansell Lab). The precision of the method expressed as RSD % was < 2%.

### Dissolved inorganic nutrients

Dissolved inorganic nitrogen compounds (NH_4_^+^, NO_2_ ^-^, NO _3_^-^) and dissolved inorganic phosphorus (PO_4_^3-^) concentrations were determined spectrophotometrically by QuAAtro segmented flow analysis (Seal Analytical, Norderstedt, Germany) following standard methods (Hansen and Koroleff, 2007). The validation and accuracy of the results were checked with reference material (Kanso Technos, Osaka, Japan) before and after sample analyses. The quality control is performed annually by participating in an intercalibration program (QUASIMEME Laboratory Performance Study).

### Dissolved amino acid analysis

Samples for total dissolved amino acid analyses were filtered through combusted GF/F filters (Whatman plc, Maidstone, UK) using acid-washed, Milli-Q water-rinsed, and combusted glass filtration system. The filtrate was collected in a dark glass vial and stored at -20°C until analysis. For each sample two technical replicates of approximately 4 mL were collected. Samples were analysed for dissolved free amino acids (DFAA) and total dissolved hydrolysable amino acids (TDHAA). For TDHAA analysis, 500 µL of the sample were first hydrolysed according to (Kaiser and Benner, 2005) with some modifications described elsewhere (Tinta et al., 2020). For DFAA analysis, 500 µL of the sample were used directly. For both measurements the samples were pipetted into acid-washed, Milli-Q wate-rinsed, and combusted glass HPLC ampules and analyzed on a Shimadzu Nexera X2 ultra high-performance liquid chromatograph (UHPLC) with a fluorescence detector (RF-20A XS; Shimadzu, Kyoto, Japan). Pre-column derivatization was applied with *ortho*-phthaldialdehyde (OPA) following (Jones et al., 1981) with slight modifications described in (Tinta et al., 2020).The concentration of dissolved combined amino acids (DCAA) was calculated as the difference between TDHAA and DFAA.

### Total bacterial abundance

To determine the total bacterial abundance, two technical replicates of 1.5 mL were fixed with 0.2 µm-filtered 37% formaldehyde (2% final concentration) and immediately stored at -80°C. Then, using a glass filtration system and a vacuum pump at low pressure (< 200 mbar), 1 mL of each replicate was filtered onto an 0.22 µm white polycarbonate filter supported by an 0.45 µm cellulose acetate filter, and stained with 2 µg mL^-1^ 4′,6-diamidino-2-phenylindole (DAPI) in Vectashield (Vector Laboratories, Newark, CA, USA). The cells were enumerated using a Zeiss Axio Imager M2 epifluorescence microscope (Carl Zeiss AG, Oberkochen, Germany) at 1000× magnification and the DAPI filter set (Ex/Em = 358/461 nm). The bacterial abundance was calculated based on the average number of cells from at least 20 counting fields with 20-200 cells enumerated per counting field.

### Bacterial cell-specific respiration and biomass production

The abundance of respiring bacterial cells was determined at a single-cell level using BacLight Redox Sensor Green Vitality Kit (ThermoFisher Scientific, Rockford, IL, USA). The Redox Sensor Green (RSG) dye produces green fluorescence (Ex/Em = 495/519 nm) when modified by bacterial reductases, many of which are part of electron transport systems and can therefore serve as a proxy for bacterial respiration (Kalyuzhnaya et al., 2008; Munson-McGee et al., 2022). From each sample, a technical triplicate of 5 mL was spiked with RSG (final concentration of 1 µM), incubated in cultivation tubes with vent caps at *in situ* temperature in the dark for 30 min, fixed with 0.2 µm filtered 37% formaldehyde (2% final concentration) and stored at -80°C. Prior to the microscopic analysis, the samples were filtered and stained with DAPI (see Total bacterial abundance section).

The biomass production of the bacterial community was determined at the single-cell level based on the incorporation rates of the methionine analog L-homopropargylglycine (HPG) into newly synthesized bacterial proteins (Samo et al., 2014). The incorporation of HPG was detected using click chemistry, where the alkyne-modified HPG is detected with Alexa Fluor 488 azide (Ex/Em = 490/525 nm), following the manufacturer’s protocol (Click-iT HPG Alexa Fluor 488 Protein Synthesis Assay Kit; ThermoFisher Scientific, Rockford, IL, USA). From each sample, a technical triplicate of 5 mL was spiked with 50 μM HPG (final concentration of 20 nM), incubated in cultivation tubes with vent caps at *in situ* temperature in the dark for 4 h, fixed with 0.2 µm filtered 37% formaldehyde (2% final concentration) and stored at -80 °C. Prior to the microscopic analysis, the samples were filtered and stained with DAPI (see Total bacterial abundance section). The filter slices were then processed according to click reaction protocols as follows: incubation in 200 μL of Click-It reaction buffer (154.5 μL Sigma water, 20 μL Click-It reaction buffer, 20 μL 10× reaction buffer additive, 4 μL copper (II) sulfate, 1.6 μL Alexa Fluor 488 azide) in the dark at room temperature for 30 min, followed by a Milli-Q water rinse and air-drying.

To quantify respiring or biomass producing cells (RSG and HPG, respectively) in specific bacterial populations, additional filter slices from both RSG and HPG incorporation procedures were used for fluorescence in situ hybridization (FISH). The filter slices were labelled with taxa-specific oligonucleotide probes (Table S4) with Cy3 at the 5’-end (Biomers, Ulm/Donau, Germany) as described in (Tinta et al., 2020).

All filter slices were analyzed using a Zeiss Axio Imager M2 epifluorescence microscope (Carl Zeiss AG, Oberkochen, Germany) at 1250× magnification, using the DAPI (Ex/Em = 358/461 nm) and the FITC (Ex/Em = 495/519 nm) filter sets, as well as the Cy3 fluorophore (Ex/Em = 554/568 nm) filter in case of FISH. At least 20 fields were counted for each filter slice using the Automated Cell Measuring and Enumeration Tool (ACMETool2, M. Zeder, Technobiology GmbH, Buchrain, Switzerland). The total abundance of respiring or biomass producing cells was determined as simultaneous signal of DAPI and FITC channels. Taxa-specific abundance of respiring or biomass producing cells was determined as simultaneous signal of DAPI, FITC, and Cy3 channels.

### Isolation and sequencing of DNA

For isolation of nucleic acids, a subsample of 0.5 L and 1 L was used from each Cteno-OM and control microcosm replicate, respectively, as well as 2L of the seawater inoculum. Bacterial biomass was collected from each sample using acid-washed, Milli-Q water-rinsed, and combusted filtration sets applying a low (<200 mbar) pressure. Total nucleic acids were extracted from the filters following (Angel et al., 2012) as modified by (Tinta et al., 2020). The extracted DNA was pooled from all Cteno-OM treatments and the control. Then, from all three DNA samples (Cteno-OM, control and inoculum) a metagenomic DNA library was prepared using a Westburg kit with enzymatic shearing (Westburg Life Sciences, Utrecht, The Netherlands) and sequencing was performed on a single lane of the HiSeqV4 Illumina platform at the Vienna Biocenter Core Facilities (https://www.viennabiocenter.org/vbcf/next-generation-sequencing/).

### Metagenomic co-assembly and analysis

The metagenomes were investigated using Anvi’o v7.0 (Eren et al., 2021) using the default parameters for each step. The reads of all three metagenomic libraries were subject to quality filtering using Fastp v0.23.2 (Chen et al., 2018). The metagenomes were co-assembled using SPAdes v3.14.1 (Prjibelski et al., 2020), followed by gene calling using PRODIGAL v2.6.3 (Hyatt et al., 2010). The identified genes were taxonomically classified using Kaiju v1.7.3 (Menzel et al., 2016) against NCBI RefSeq database from 26.02.2021 (Pruitt et al., 2005). Functional annotation of the genes was carried out using DIAMOND v2.0.15.153 (Buchfink et al., 2021) against the NCBI Clusters of Orthologous Groups of proteins (COG) database (Tatusov et al., 2000), and using KofamKOALA against the KEGG KOfam database (Aramaki et al., 2020).

The reads of each library were mapped to the co-assembled metagenome using BBmap v37.61 (sourceforge.net/projects/bbmap/), part of BBTools (Bushnell et al., 2017). Then the metagenome was binned using CONCOCT v1.1.0 (Alneberg et al., 2014) and MetaBAT v2.16 (Kang et al., 2019) and then merged using DAStool v1.1.6 (Sieber et al., 2018). The resulting metagenome-assembled genomes (MAGs) were further refined (Eren et al., 2015). The completeness of metabolic KEGG modules (Kanehisa, 2017; Kanehisa et al., 2014) in each MAG was estimated using the anvio program ‘anvi-estimate-metabolism’ (see tutorial: https://merenlab.org/m/anvi-estimate-metabolism).

For determining the exact phylogeny of the selected MAG, complete genomes of *Pseudoalteromonas* were retrieved from NCBI genome database. The phylogenetic tree was generated using trimAl v1.4.rev15 (Capella-Gutiérrez et al., 2009) and IQ-TREE v2.2.0.3 (Nguyen et al., 2015), based on single copy genes identified using hidden Markov model search in each genome.

The selected *P*.*phenolica* genomes (Table S5) were then annotated as described above and the pangenome was reconstructed using anvio program ‘anvi-pan-genome’ (Delmont and Eren, 2018) with MCL hierarchical clustering of the genes (van Dongen and Abreu-Goodger, 2012). The average nucleotide identity was calculated using PyANI v0.2.12 (Pritchard et al., 2015).

### Isolation and mass spectrometry analysis of proteins

For proteomic analysis, extraction of soluble proteins from ctenophore biomass and from treatments’ media (i.e., exo-metaproteomes) was performed as described in detail in Tinta et al. (2020). For analyses of endo-metaproteomes, biomass was collected onto 0.22 µm polycarbonate filters. For the coastal endo-metaproteome 3 L of the microbial inoculum were collected prior to the start of the experiment; 300 mL were collected from each ctenophore treatment and 1 L from each control treatment at the peak of the bacterial abundance. Protein extraction from collected cells was performed as follows: filters were ground into small pieces with a sterile metal spatula after submerging the tubes with the filters into liquid nitrogen. Filter pieces were resuspended in lysis buffer (100mM Tris-HCl pH 7.4, 1% SDS, 150mM NaCl, 1mM DTT, 10mM EDTA) and cells were lysed with five freeze-and-thaw cycles. After centrifugation (20,000 g at 4°C for 25 min) the supernatant was transferred into a tube and proteins were co-precipitated with 0.015% deoxycholate and 7% trichloroacetic acid (TCA) on ice for 1h and washed twice with ice-cold acetone. Dried protein pellets were resuspended with 50 mM TEAB buffer (Millipore Sigma, Burlington, MA, USA) and quantified using Pierce 660nm Protein Assay Reagent (ThermoFisher Scientific, Rockford, IL, USA). Next, cysteines were reduced and alkylated with 10 mM DTT and 55 mM iodoacetamide (IAA), respectively.

For analyses of exo-metaproteomes, the filtrate (< 0.22 µm) of each sample was concentrated using a VivaFlow 200 (Sartorius, Göttingen, Germany) with 30 KDa and 5 KDa Molecular Weight Cut-Off (MWCO) to collect the high molecular weight (30 KDa - 0.22 μm) and the low molecular weight (5 KDa - 30 KDa) fraction, respectively. The high and low molecular-weight fractions were further concentrated to 250 μL using an Amicon (Millipore Sigma, Burlington, MA, USA) Ultra-15 Centrifugal Filter 30 KDa MWCO and 3 KDa MWCO Unit. Sample reducing agent NuPAGE (Invitrogen, Waltham, MA, USA) was added to the samples to reach 1X final concentration.

All samples (endo- and exo-metaproteomes) were re-precipitated using 9 times the sample volume of 96% EtOH at -20°C overnight. Pellets were resuspended in 50 mM TEAB, followed by overnight in-solution trypsin (Roche, Basel, Switzerland) digestion (1:100, w/w) at 37°C. TFA was added to the samples at 1% final concentration to terminate trypsin digestion. Samples were desalted using Pierce C18 Tips (ThermoFisher Scientific, Rockford, IL, USA) according to the manufacturer’s protocol. Prior the LC MS/MS analyses, digested peptides were dissolved in 0.1% formic acid and 2% acetonitrile and transferred into micro-inserts sealed with aluminum caps. Prior to the run, the concentration of peptides was measured using Pierce Quantitative fluorometric peptide assay (ThermoFisher Scientific, Rockford, IL, USA).The resulting peptides were sequenced on a Q-Exactive Hybrid Quadrupole-Orbitrap Mass Spectrometer (ThermoFisher Scientific, Rockford, IL, USA) at the Vienna Research Platform for Metabolomics & Proteomics and analyzed using the Proteome Discoverer v2.2.0.388 (ThermoFisher Scientific, Rockford, IL, USA) at the Life Science Computer Cluster (LiSC) of the University of Vienna. The tandem mass spectrometry spectra of proteins extracted from the Cteno-OM powder were searched using MASCOT v2.6.1 (Perkins et al., 1999) against the *M. leidyi* transcriptome shotgun assembly project (NCBI TSA accession number GFAT01000000). The tandem mass spectrometry spectra of proteins extracted from the microcosms and the seawater inoculum were searched using SEQUEST-HT against the bacterial protein coding genes of the co-assembled metagenome. Search parameters were as follows: enzyme - trypsin, fragment mass tolerance - 0.8 Da, max. missed cleavages - 2, fixed modifications - carbamidomethyl (Cys), optional modifications - oxidation (Met). Percolator parameters were as follows: max. delta Cn: 0.6, max. rank: 0, validation based on q-value, false discovery rate (calculated by automatic decoy searches) 0.05. Protein quantification was conducted using the chromatographic peak area-based label-free quantitative method.

### Statistical analyses

Statistical analyses were done in R v4.2.1 (R Core Team 2022) using RStudio v2022.02.3 (RStudio Team 2019). The metagenomic and metaproteomic datasets were combined and managed using ‘phyloseq’ v1.40 (McMurdie and Holmes 2013). Dissimilarities between samples and statistical tests were carried out using ‘vegan’ v2.6-2 (Oksanen *et al*. 2022). Protein enrichment analysis was performed using ‘DESeq2’ v1.38.3 (Love et al., 2014) on a variance-stabilized protein abundance matrix. To avoid overrepresentation, per protein only the COG assignment with the highest e-value was considered in the summary of the results. Mapping of the genes to KEGG pathways was carried out using ‘pathview’ v1.38.0 (Luo and Brouwer, 2013).

### Data availability

The metagenomic raw sequences have been deposited in the European Nucleotide Archive (ENA) at EMBL-EBI under Project accession number PRJEB63998. The mass spectrometry proteomics data have been deposited to the ProteomeXchange Consortium (Deutsch et al., 2023) via the PRIDE (Perez-Riverol et al., 2022) partner repository with the dataset identifier PXD043478.

Scripts for the molecular data processing and statistical analyses can be accessed via Zenodo (https://www.zenodo.org/) under doi: 10.5281/zenodo.8119924.

## Author contributions

EF performed the bioinformatic and statistical analyses and wrote the manuscript. TT and GJH designed the experiments. TT and JHH conducted the experiments and performed laboratory analyses. KK performed chemical analysis of inorganic nutrients and dissolved organic matter. ZZ contributed and assisted with experimental design and preparation of samples for proteomic analysis. CA contributed and assisted with preparation of samples for microscopy-based analysis. All authors contributed to the article and approved the submitted version.

## Acknowledgments

We thank Sonja Tischler for performing mass spectrometry. We thank Barbara Mähnert for help with experimental setup and the staff of Marine Biology Station Piran and crew of RV *Sagita* for their help with sampling.

This project received funding from the European Union’s Horizon 2020 Research and Innovation Program under the Marie Skłodowska-Curie Grant Agreement No. 793778. EF and GJH were funded by the Austrian Science Fund (FWF) project I04978. TT was further supported by the Slovenian Research Agency under grant number ARRS J7-2599 and by the Slovenian Research Agency (Research Core Funding No. P1-0237).

